# Inter-subject representational similarity analysis reveals individual variations in affective experience when watching erotic movies

**DOI:** 10.1101/726570

**Authors:** Pin-Hao A. Chen, Eshin Jolly, Jin Hyun Cheong, Luke J. Chang

## Abstract

We spend much of our life pursuing or avoiding affective experiences. However, surprisingly little is known about how these experiences are represented in the brain and if they are shared across individuals. Here, we explore variations in the construction of an affective experience during a naturalistic viewing paradigm based on subjective preferences in sociosexual desire and self-control using intersubject representational similarity analysis (IS-RSA). We found that when watching erotic movies, intersubject variations in sociosexual desire preferences of 26 heterosexual males were associated with similarly structured fluctuations in the cortico-striatal reward, default mode, and mentalizing networks. In contrast, variations in the self-control preferences were associated with shared dynamics in the fronto-parietal executive control and cingulo-insula salience networks. Importantly, these results were specific to the affective experience, as we did not observe any relationship with variation in preferences when individuals watched neutral movies. Moreover, these results appear to require multivariate representations of preferences as we did not observe any significant results using single summary scores. Our findings demonstrate that multidimensional variations in individual preferences can be used to uncover unique dimensions of an affective experience, and that IS-RSA can provide new insights into the neural processes underlying psychological experiences elicited through naturalistic experimental designs.

## Introduction

Emotions play a critical role in our lives, from organizing responses relevant for our survival (Davidson and Irwin, 1999) to facilitating social connection (FeldmanHall and Chang, 2018). Though much work has sought to understand the sensory and cognitive processes involved in reliably perceiving emotional signals in pictures of facial expressions (Kim et al., 2011; Whalen et al., 2013) or arousing scenes (Chikazoe et al., 2014; Lindquist et al., 2012), considerably less is known about how the brain constructs emotional experiences (Chang et al., 2015). Emotional experiences arise from integrating exogenous information from the external world with endogenous information reflecting past experiences, current homeostatic states, and future goals (Chang et al. 2018). At perhaps the simplest level, an affective experience requires making an appraisal of whether an external stimulus is good or bad (Ashar et al., 2017; Ellsworth and Scherer, 2003). Since appraisals are cognitive evaluations of events and situations, these evaluations are not simple perceptions but rather are constructed interpretations of unfolding events (Scherer, 2013, 1999). As stimuli do not inherently contain value, these judgments are often made with respect to specific goals (e.g., I don’t want to embarrass myself) (Chang and Jolly, 2018; FeldmanHall and Chang, 2018), homeostatic needs (e.g., I’m hungry) (Panksepp, 2004), and interpersonal judgments and comparisons (e.g., am I doing as well as this other person?) (Bault et al., 2011; Kumaran et al., 2016). For example, certain types of foods that might be aversive, such as salt to a rodent, instantly become much more appealing when the animal becomes deficient in sodium (Robinson and Berridge, 2013). This variation in endogenous states makes studying affective experiences extraordinarily difficult as different individuals might experience the same stimuli differently and the same individual’s experience may change over repeated exposures (Barrett, 2006).

This complexity raises the question, what are the most appropriate dimensions to describe distinct affective experiences? This measurement problem has vexed psychologists for many decades (Larsen and Fredrickson, 1999). Should affective experiences be described using specific feelings, which can be verbally articulated or represented in configurations of facial expressions (Ekman, 1992)? The act of labeling feelings requires a conceptual representation, which may impact the experience itself (Lindquist et al., 2015; Nisbett and Wilson, 1977; Satpute et al., 2016). Alternatively, affective experiences may be better represented in a lower dimensional space such as valence and arousal (Mattek et al., 2017; Russell, 1980). However, it is hard to imagine that the entirety of the human affective experience space can be spanned by 2 or 5 dimensional representations (Jolly and Chang, 2019). This suspicion has led other researchers to use data-driven methods to find evidence for higher dimensional spaces (e.g., 27 dimensions) (Cowen and Keltner, 2017). Given these limitations, how should we study experiences? Does it even make sense to try and represent affective experiences in a space that requires language and concepts?

One possibility is to assume that affective experiences might require specific computations in distinct regions of the brain. This would imply that variations in experience might be measured by different patterns of brain activity. Prior work has shown that different affective experiences such as feeling bad from viewing pictures (Chang et al., 2015), experiencing thermal pain applied to a limb (Wager et al., 2013), or viewing others in pain (Krishnan et al., 2016) are reflected in reliable and distinct patterns of brain activity measured using functional magnetic resonance imaging (fMRI). For each of these affective experiences, the intensity of the experience can be reliably predicted in most participants by using a common pattern of brain activity (Woo et al., 2017). However, if affective experiences vary across individuals, these variations would also presumably be reflected in patterns of brain activity (Chang et al. 2018), which would not be adequately described by a single common model. Though recent developments in functional alignment techniques have shown promise in developing models that better generalize across people (Chen et al., 2015; Guntupalli et al., 2016; Haxby et al., 2011), these techniques assume that participants have identical functional responses and may be suboptimal when aligning on individual affective experiences that may have unique temporal dynamics (Chang et al. 2018). Thus, identifying regions that vary in the experience may provide important insights into characterizing what types of psychological and neural processes are involved in constructing the affective experience itself.

To test this hypothesis, we explored how different aspects of an affective experience might vary across individuals based on individual differences. Specifically, we observed the neural dynamics of heterosexual male college undergraduates (n=26) while viewing short clips of erotic and neutral movies using fMRI (Figure 1). We hypothesized that this type of naturalistic stimuli and viewing environment would create a unique affective experience. On the one hand, this particular population might enjoy viewing erotic movies, and individuals with similar sociosexual desire preferences might show more similar patterns of brain responses in regions involved in the cortico-striatal reward and mentalizing networks (Ferretti et al., 2005; Gillath and Canterberry, 2012; Walter et al., 2008) compared to those with different preferences. On the other hand, viewing erotic movies in this particular neuroimaging environment while being monitored by research staff might also introduce a level of self-consciousness, leading to increased self-control thereby changing indivduals’ experiences. We also explored how individual variations in both subjective and task-derived objective self-control might impact the experience reflected through variations in the fronto-parietal executive control and cingulo-insula salience networks (Aron, 2011; Aron et al., 2007; Heatherton and Wagner, 2011). We predicted that individual variations in these two preferences would specifically be involved in the generation of the affective experience when individuals were watching erotic movies. However, we did not expect that these individual variations would reflect general individual differences, which could be observed in any type of experience such as viewing neutral movies. We also assumed that these preferences would be complex and that individual variations would be better captured in a multidimensional feature space then a single summary score, where each item from the questionnaires would serve as an independent dimension. Thus, we used intersubject-representational similarity analysis (IS-RSA) (van Baar et al., 2019) to explore how individual differences in sociosexual desire and self-control preferences impacted the inter-indivdual variation in the experience of viewing erotic compared to neutral movies.

**Figure 1.**
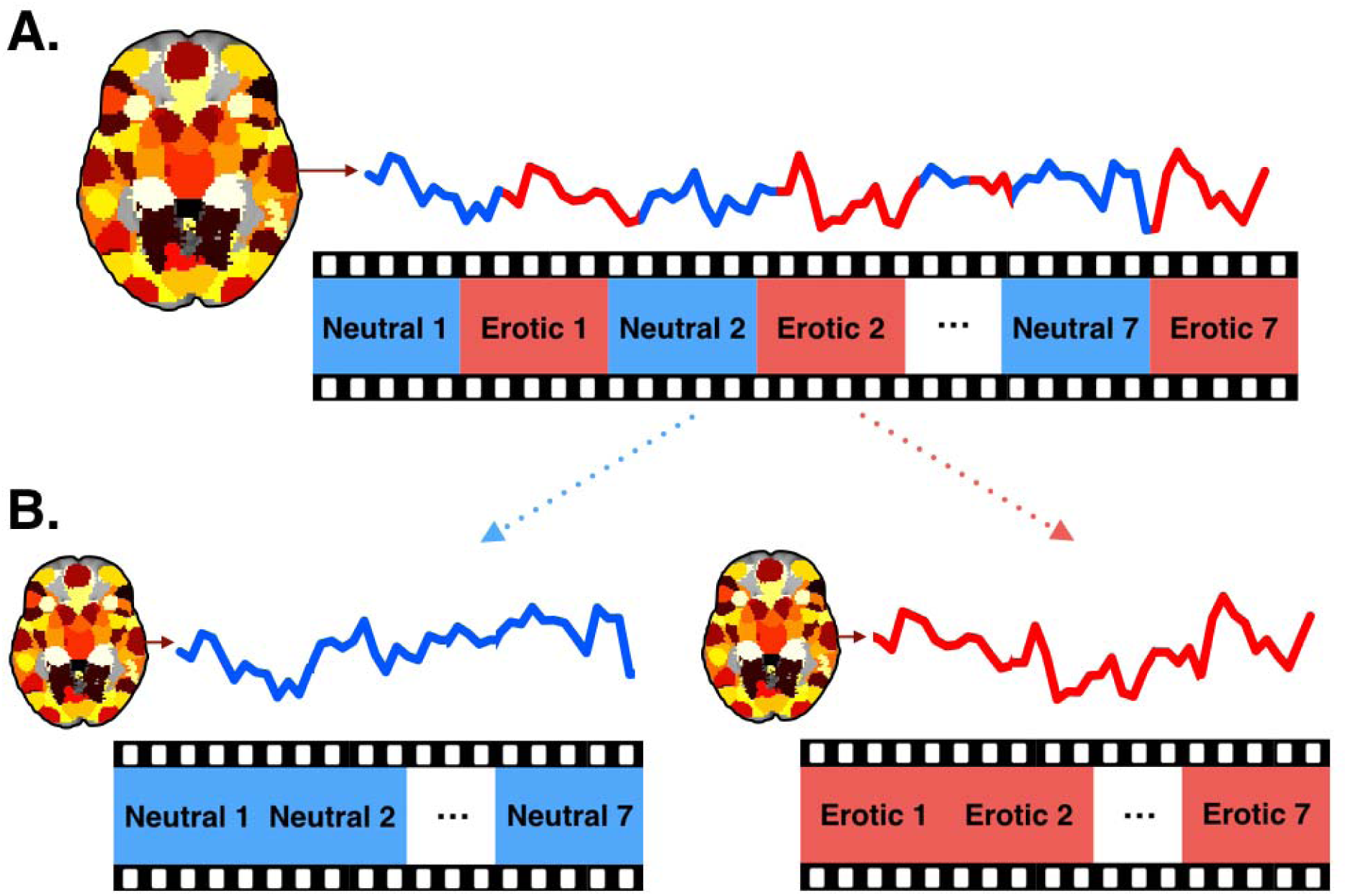
Naturalistic viewing task and extraction of time course data. (A) Participants watched a 7-minute long movie containing 7 short neutral movies and 7 erotic movies in the fMRI scanner, and the sequence of movies is identical to each participant. (B) For each participant, time courses for the neutral-movie condition and erotic-movie condition were concatenated and extracted from 100 non-overlapping brain ROIs.

## Methods

### Participants

Twenty-six right-handed heterosexual male undergraduates (age range: 18-26; 76% Caucasian, 17% Asian, 7% Hispanic or other) were recruited from Dartmouth College. No participants had a history of mental health or neurological disorders. Participants gave informed consent in accordance with the guidelines set by the Committee for the Protection of Human Subjects at Dartmouth College.

### Naturalistic viewing task

Participants were asked to watch a 7-minute long compilation of video clips while undergoing functional magnetic resonance imaging (fMRI). This 7-minute movie contained 7 short neutral clips as well as 7 short softcore erotic clips. Each clip lasted for 30 seconds and alternated between the neutral and erotic videos. The sequence of the movies first began with a neutral movie, followed by an erotic movie, and ended with the seventh erotic movie (Figure 1A). Most importantly, each participant watched the same movies in the same sequence, ensuring that the temporal order input was matched across participants during scanning. These 7 short erotic movies were selected from a large pool of erotic movies. This selection process was performed by two research assistants, and the selection criteria was based on whether both rated a particular clip a 5, the highest rating, on a 5-point sexual arousal scale. The 7 short neutral movies were selected from a large pool of clips downloaded from an online sharing platform. These neutral movies were related to natural scenes, including forest, hills, plains, beaches and the sea. Videos were selected among clips that were rated by both research assistants the lowest rating on the sexual arousal scale.

### Behavioral measurements

#### Sociosexual desire

To understand how participants perceived their sociosexual desire and experience, participants completed two widely-used questionnaires after the scanning session: the revised Sociosexual Orientation Inventory (SOI-R; Penke and Asendorpf, 2008), and Sexual Desire Inventory-2 (SDI-2 Spector et al., 1996). The SOI-R contains nine items and covers three different dimensions, including past sexual experiences, the attitude toward casual sex, and sociosexual desire (e.g. how often do you experience sexual arousal when you are in contact with someone you are not in a committed romantic relationship with?). The SDI-2 contains 14 items focusing on sexual desire, including solitary and dyadic sexual desire (e.g. during the last month, how often have you had sexual thoughts involving a partner?). Together, these two questionnaires contain 23 behavioral items and cover several distinct aspects of the subjective experience of sociosexual desire.

#### Self-control

In order to examine how participants subjectively perceived their self-control experiences, they completed the brief self-control scale (Tangney et al., 2004), a widely-used 13-item questionnaire in self-control research. These items covered several distinct domains, including impulse control, thought suppression, behavioral inhibition, and goal achieving. In addition to these subjective measurements, we also included a Go/No-go task as an objective measure of self-control. Prior to completing the task, each participant selected the 18 most attractive pictures from a set of 45 attractive female pictures, and another 18 pictures that were perceived to be the least attractive from a separate set of 45 unattractive pictures, which were based on an attractive norm from a group of 6 independent heterosexual male raters. These 18 attractive and unattractive pictures were used as stimuli in the Go/No-go task. There were four runs in this task, and the rule switched across runs. For example, the rule was to “Go” for the attractive targets and “No go” for the unattractive targets in the first run, but the rule reversed to “Go” for the unattractive and “No go” for the attractive targets in the second run. For each trial, the picture was shown for 500 milliseconds followed by a fixation for 2000 milliseconds. Participants completed 4 runs, which included 54 Go trials and 18 No-go trials for each condition per run. Accuracy was calculated by computing the average number of correctly inhibited responses in the No-go trials and correct button responses during the Go trials separately for each condition, resulting in four indices to represent an objective measure of self-control. Thus, by using both subjective and objective measurements, there were 17 behavioral features capturing individual variations in self-control preferences.

#### Imaging

Participants were scanned using a Philips Intera Achieva 3T scanner with a 32 channel head coil. Functional images were acquired using a T2*-weighted echo-planar sequence (TR = 2500 ms, TE = 35 ms, 90 flip angle, FOV = 240 mm, 36 axial slices, 3 mm thick with 0.5 mm gap, 3 × 3 mm in-plane resolution). Structural images were acquired by using a T1-weighted MPRAGE sequence (160 sagittal slices, TR = 9.9 ms, TE = 4.6 ms, 8 flip angles, 1 × 1 × 1 mm voxels).

#### fMRI data preprocessing

Imaging data were preprocessed using SPM8 (Wellcome Department of Cognitive Neurology, London, England) in conjunction with a toolbox for preprocessing and analysis (available at http://github.com/ddwagner/SPM8w) and the nltools package 0.3.14 (Chang et al., 2018). Within each functional run, differences in acquisition time between slices were first corrected. Images were then realigned within and across functional runs using affine registration with 6 degrees of freedom for motion correction. After motion correction, images were normalized with a 12-parameter affine transformation and non-linear transformation into a standard stereotaxic space (3 ×3 × 3mm isotropic voxels), the SPM8 EPI template, which conforms to the ICBM 152 brain template space. After normalization, a 6-mm full-width-at-half-maximum Gaussian kernel was applied for spatial smoothing. Lastly, for each participant, covariates of no interest (a session mean, a linear trend, six movement parameters, the square of the six movement parameters, the temporal derivative of the six movement parameters, and the square of the derivatives) were regressed out from the smoothed images using ordinary least squares regression.

#### Inter-subject correlation

We first used intersubject correlation (ISC) to examine the reliability of neural dynamics responding to a dynamic stimuli across individuals (Cohen et al., 2017; Hasson et al., 2004; Nastase et al., 2019). To perform this analysis, for each participant we extracted the timepoints when participants were watching the erotic and neutral movies and separately concatenated all of the videos into two separate conditions. We then separately extracted the mean time course for each condition with 100 non-overlapping regions defined from a whole-brain parcellation that divided the brain into 100 functionally similar regions-of-interest (ROIs) based on meta-analytic functional coactivation of the Neurosynth database (de la Vega et al., 2016) (Figure 1B; parcellations available at http://neurovault.org/media/images/2099/). For each stimulus type, we separately computed the pairwise correlation between participants’ mean time-course in each ROI producing a subject by subject correlation matrix for each ROI. (Figure 2A(1)). To compare the spatial of mean ISC between conditions, we computed the mean of the lower triangle of the subject by subject correlation matrix with each ROI and then correlated these mean values between erotic and neutral movies (Figure 3).

**Figure 2.**
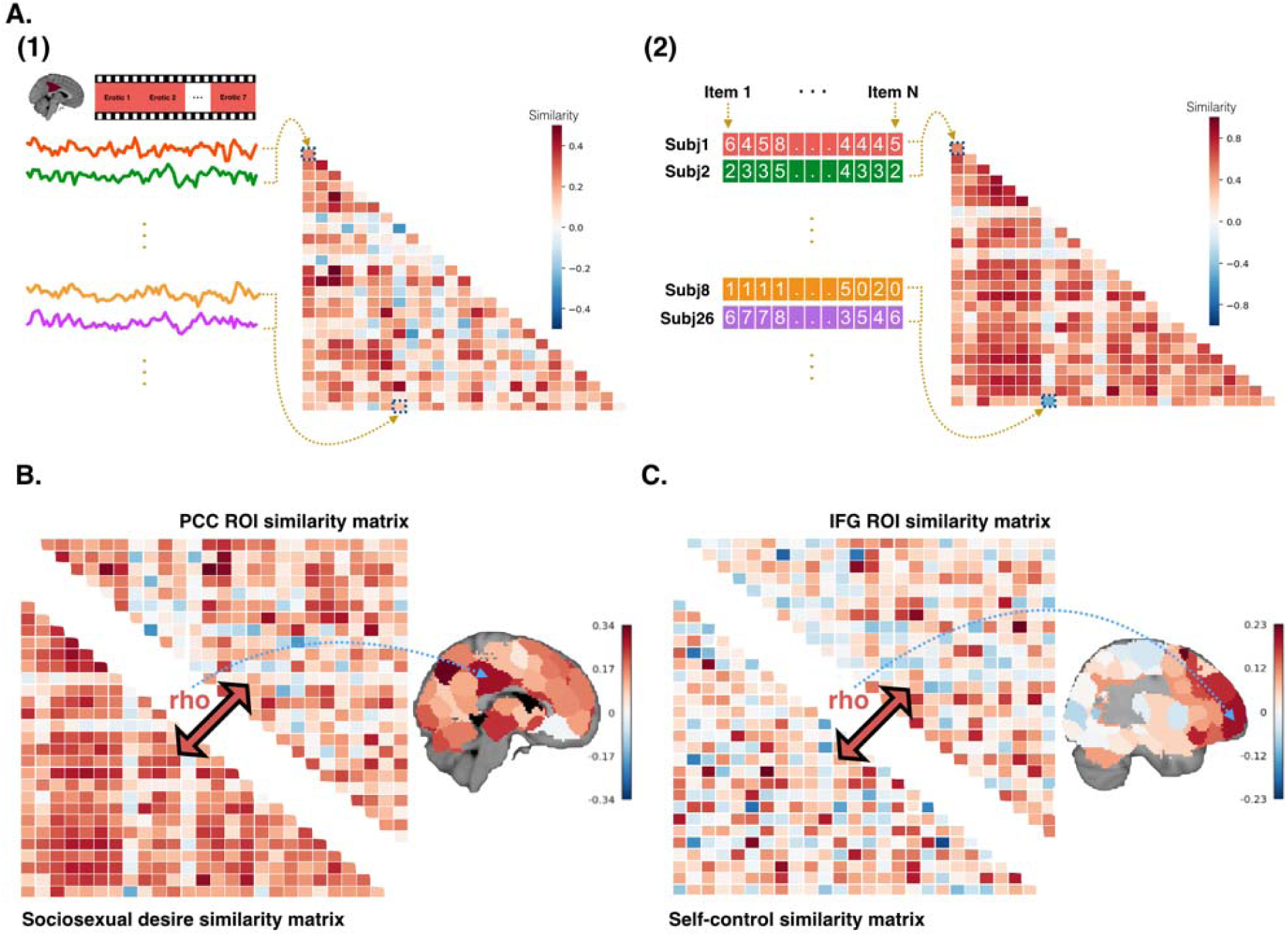
Intersubject-representational similarity analysis. (A) IS-RSA involves first computing (1) a subject by subject correlation matrix using mean the time-series responses within an ROI for erotic movies and (2) a subject by subject correlation matrix using item behavioral item responses for sociosexual desire preferences. Each matrix reflects inter-subject variability in brain responses and preferences respectively. (B) We find that inter-subject variability in PCC is captured well by inter-subject variability in sociosexual desire using spearman rank correlation. (C) Similarly, we find that inter-subject variability in IFG is well captured by inter-subject variability in self-control.

**Figure 3.**
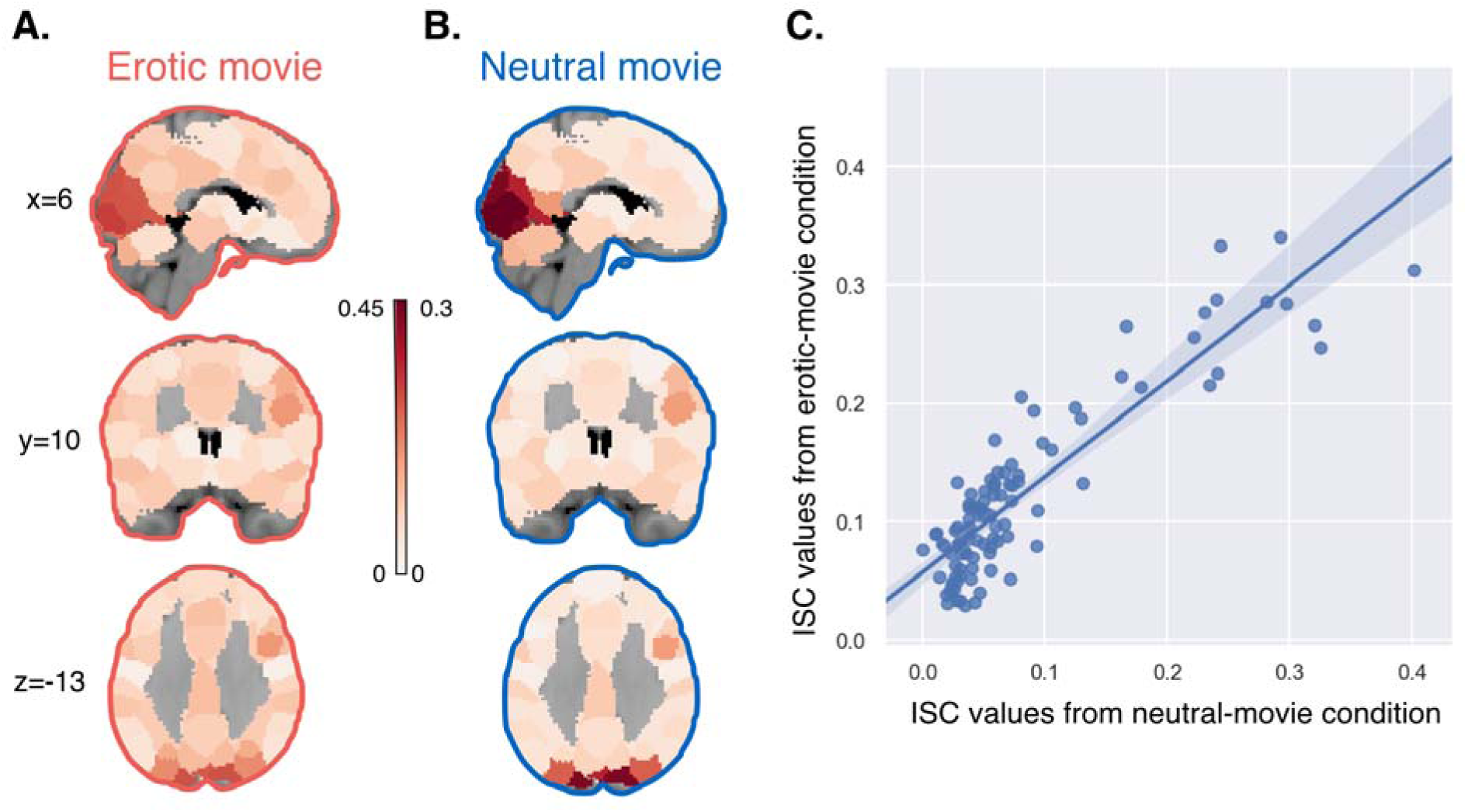
Intersubject correlation. Mean ISC within each ROI separately for (A) the erotic-movie condition and (B) the neutral-movie condition. The color bar indicates the average ISC value within each ROI for each condition respectively. (C) The spatial distribution of these maps was highly similar between the two conditions (each point reflects a single ROI).

#### Inter-subject representational similarity analysis

We used inter-subject representational similarity analysis (IS-RSA) (van Baar et al., 2019) to identify regions of the brain that were involved in processing distinct aspects of participant’s affective experience. This analytic technique is an extension of intersubject correlation analysis (ISC), which identifies the reliability of temporal responses to a dynamic stimuli across participants (Cohen et al., 2017; Hasson et al., 2004). However, rather than examining the reliability of responses in a region across participants, IS-RSA instead explores how inter-individual variability in brain dynamics is related to individual differences in behavioral disposition using second order statistics akin to representational similarity analysis (RSA) (Kriegeskorte et al., 2008; Nguyen et al., 2019; Nummenmaa et al., 2012). The intuition is that participants who occupy a similar position in the multidimensional feature space (e.g., sociosexual desire or self-control) will be processing information about the video more similarly and regions involved in these processes should show a commensurate similarity in temporal dynamics (van Baar et al., 2019).

Just like the neural ISC analyses described above, we computed the correlation between pairs of participants’ behavioral scores (Figure 2A(2)). We did this separately for each behavioral domain (i.e. sociosexual desire and self-control). To ensure that no single behavioral item could drive the similarity between participants, we normalized each behavioral feature to range [0,1] by dividing each feature with its maximal score on the scale prior to computing pairwise correlations between participants.

Finally, we used spearman rank correlations to compute the overall similarity between the lower triangles of the neural and behavioral inter-subject similarity matrices separately for each ROI (Figure 2B). This procedure yielded two separate whole-brain maps of regional similarity in individual variations in experience based on variations in sociosexual desire (Figure 2B) and self-control preferences (Figure 2C) for the erotic video-watching condition, and also yielded another two whole-brain maps for the neutral video-watching condition. Maps were thresholded using a Mantel permutation test (Nummenmaa et al., 2012), in which both the rows and columns of one subject by subject correlation matrix were shuffled and the spearman correlation between both correlation matrices was recomputed. This procedure was repeated 10,000 times to generate a null distribution of rank correlations which was used to compute p-values based on one-tailed test of correlations greater than 0 for each ROI (Nili et al., 2014). Permuted p-values were thresholded using a false-discovery rate, q < 0.05 across ROIs.

#### Meta-analytic decoding with Neurosynth

In order to decode the brain maps representing regional similarity in individual variations in experience based on variations in each behavioral domain, we used the meta-analytic decoding method to make reverse inferences of mental states from brain activation maps (Chang et al., 2013). Reverse inference reveals the degree of a particular psychological process presented by giving a specific brain activation map, which relies on a comprehensive dataset of term-to-activation mappings provided by the Neurosynth framework (http://neurosynth.org; (Yarkoni et al., 2011)). We used topic-based maps from previous studies (Fox et al., n.d.; Sul et al., 2017), in which 80 topics were generated using Latent Dirichlet Allocation topic modeling of 9,204 fMRI articles. Among the 80 topics, we selected 8 topics that are most relevant to sociosexual desire and self-control, and 8 meta-analytic reverse inference maps corresponding to the 8 topics were generated based on the mapping between each topic and brain activation at each voxel in the Neurosynth dataset. We then computed the pearson correlations between our IS-RSA brain maps and each of the meta-analytic reverse inference maps. Lastly, we used polar plots to visualize the strength of association between each topic and each of our brain maps representing regional similarity in individual variations in experience based on variations in a given behavioral domain.

## Results

### Inter-subject Correlation

We first examined the overall similarity in temporal responses in each region separately for each condition using intersubject correlation analysis (ISC). Overall, we observed slightly higher ISC when participants viewed erotic movie clips (mean ISC = .12), compared to viewing the neutral clips (mean ISC = .08), *t(99)* = 11.35, *p* = 0.002. We found that the highest average ISC values were observed in ROIs within the occipital lobe and ventral temporal lobe (Figure 3A & 3B), and that the spatial distribution of ISC values across the whole brain was highly consistent across the two conditions, *r* = .89, *p* < .001 (Figure 3C). This suggests that though the erotic movie clips might have produced a slightly higher degree of alignment of temporal brain responses, the overall spatial topography of ISC was highly similar when viewing videos from the two conditions.

### Intersubject Representational Similarity Analysis

Next, we performed intersubject representational similarity analysis (IS-RSA) to identify brain regions with temporal dynamics that showed similar patterns of inter-individual variability to either sociosexual desire or self-control preferences. Importantly, inter-individual variation in sociosexual desire was not related to variation in self-control, *r* = -.01, *p* = .89, suggesting that both measures captured different aspects of variability across individuals. Variation in sociosexual desire was postively reflected in brain regions within the cortico-striatal reward and default mode as well as mentalizing networks when participants were watching erotic movies (Figure 4A). In particular, the highest similarities were found in the nucleus accumbens (NAcc; *r* = .23, *p* = 0.002), precuneus (*r* = .34, *p* = 0.0002), posterior cingulate cortex (PCC; *r* = .28, *p* = 0.001), midbrain (*r* = .24, *p* = 0.0017), middle temporal gyrus (*r* = .24, *p* = 0.001), temporoparietal junction (TPJ; *r* = .28, *p* = 0.0004), and the middle frontal gyrus (*r* = .26, *p* = 0.002) (Figure 6A). Importantly, these results were specific to the affective experience of watching erotic movies as no such similarity was found within these regions when participants watched neutral movies (Figure 4B). Instead, variations in sociosexual desire were negatively correlated with variations in brain responses within the medial prefrontal cortex (MPFC) and the precuneus.

**Figure 4.**
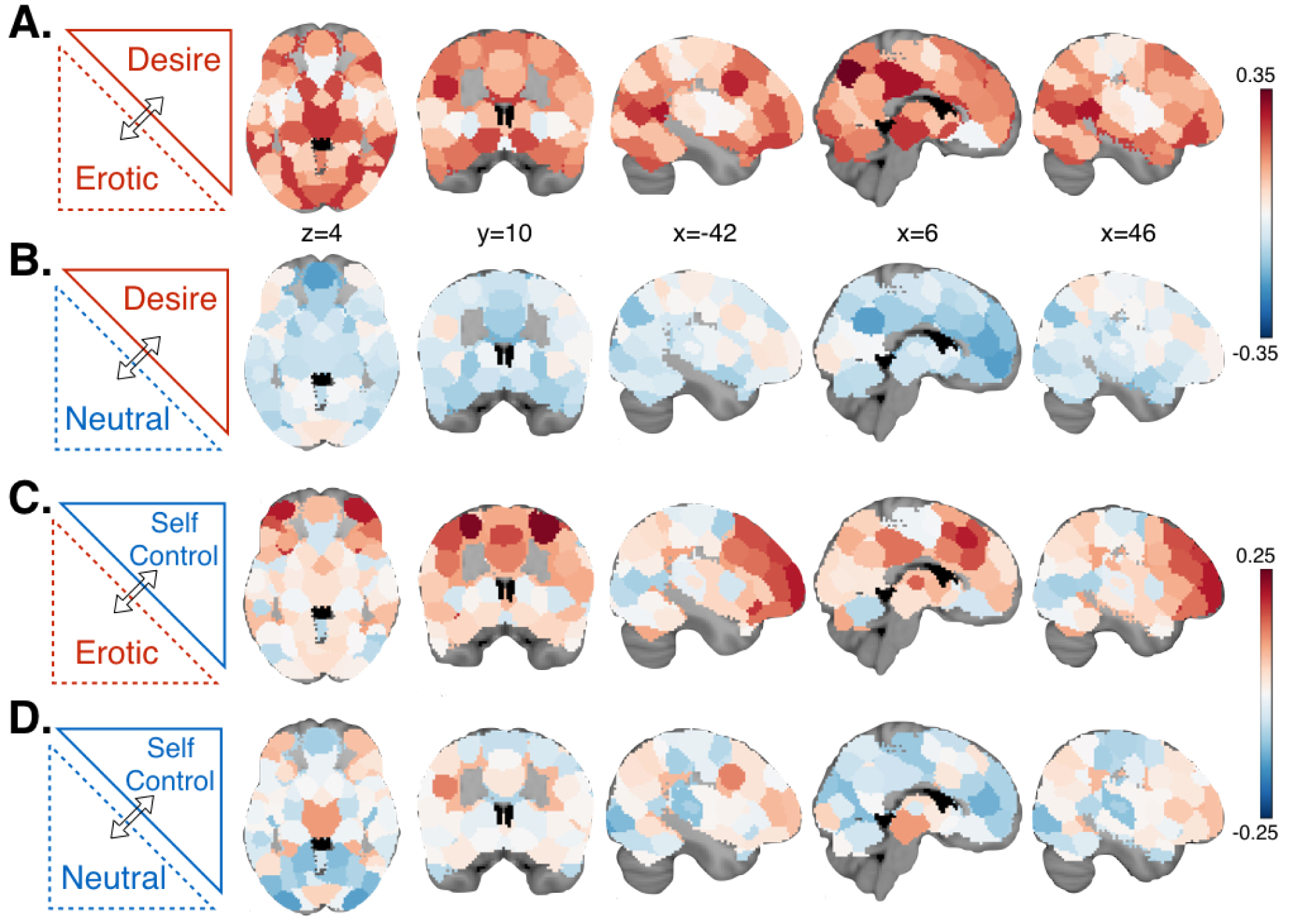
Whole-brain IS-RSA un-thresholded similarity maps. (A) Individual variations in experience based on variations in sociosexual desire showed higher similarity in the cortico-striatal reward and default mode as well as mentalizing networks when watching erotic movie, (B) but no such similarity was shown when watching neutral movies. (C) In contrast, variations in experience based on variation in self-control showed higher similarity in the fronto-parietal executive control and cingulo-insula salience networks when watching erotic movies, (D) but no such similarity was shown when watching neutral movies. The color bar indicates the value of spearman correlation rho for each ROI from each IS-RSA analysis.

**Figure 5.**
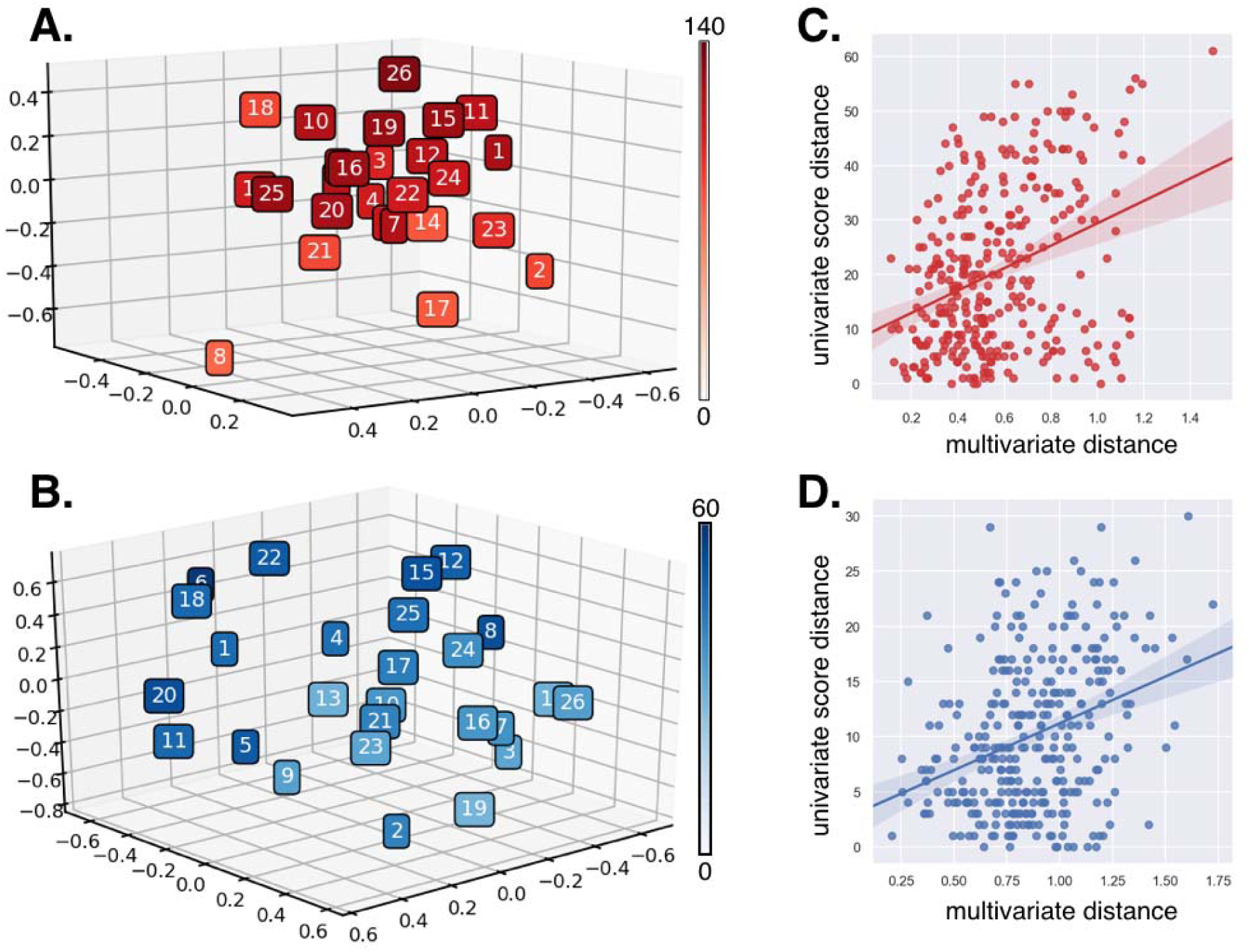
Multivariate intersubject similarity structures. The relative position of each participant in a multivariate space for their (A) sociosexual desire and (B) self-control preferences. The darker color on the color bar indicates a higher univariate summary score for each behavioral domain. The spearman correlation between the multivariate and univariate summary score distance matrices for (C) the sociosexual desire and (D) self-control preferences.

**Figure 6.**
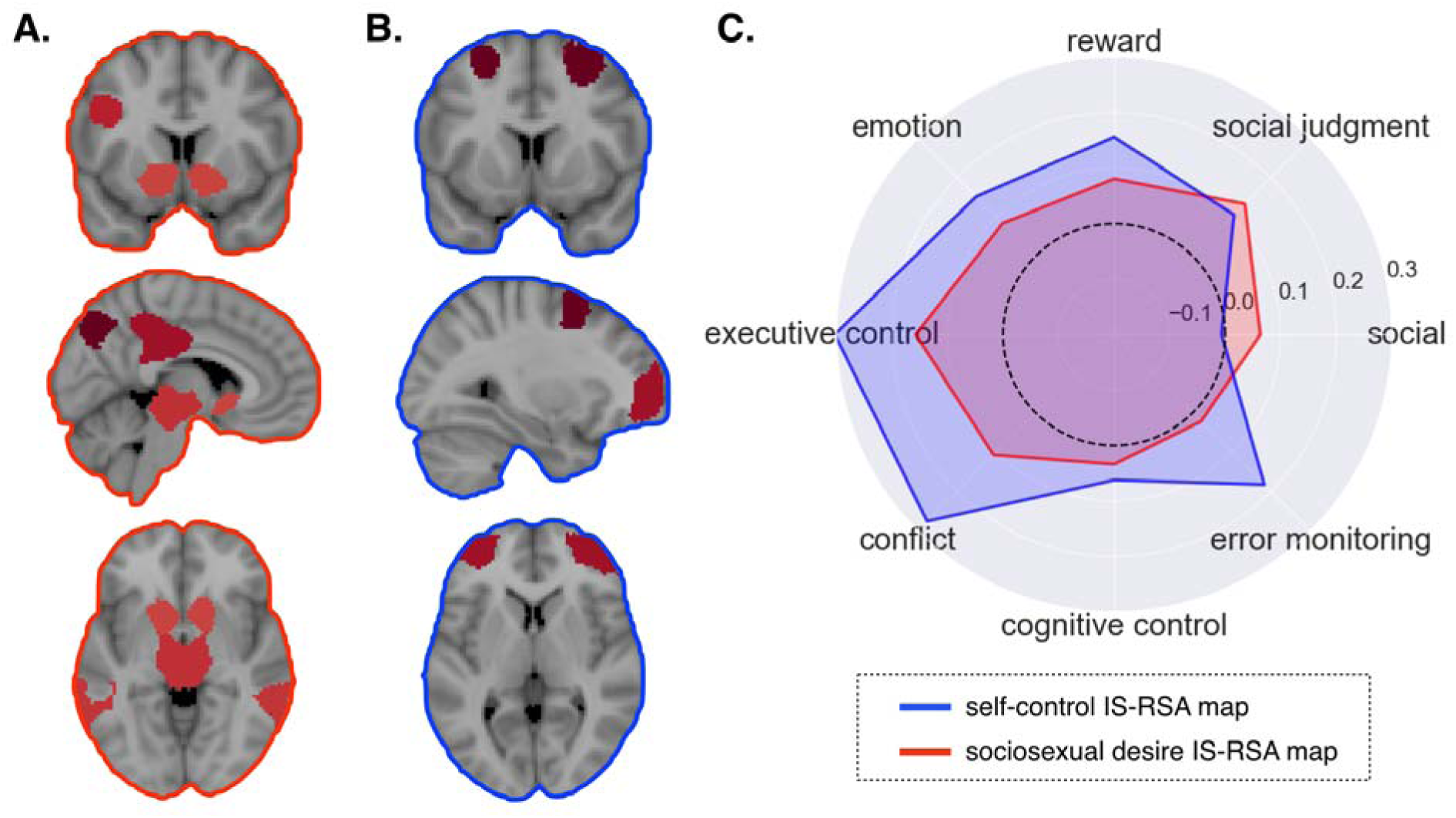
Thresholded IS-RSA maps and meta-analytic decoding. Thresholded inter-subject similarity map based on variations in (A) the sociosexual desire and in (B) the self-control preferences. (C) The unthresholded map based on sociosexual desire revealed stronger association with social and social judgment, whereas the thresholded similarity map based on self-control revealed stronger association with emotion, reward, executive control, cognitive control, error monitoring, and conflict.

Next, we examined where inter-individual variations in self-control captured variation in neural dynamics. Variations in self-control were positively associated with variations in brain regions within the fronto-parietal executive control and cingulo-insular salience networks when participants were watching erotic movies (Figure 4C). In particular, the highest similarities were found in the superior frontal gyrus (SFG; *r* = 0.23, *p* = 0.0003) and the inferior frontal gyrus (IFG; *r* = .19, *p* = 0.0006) (Figure 6B). In contrast, when participants were watching neutral movies, no significant similarity was found within these two networks (Figure 4D).

### Feature space dimensionality

We predicted that individual variations in preferences would be better captured using a multidimensional representation compared to the more traditional use of single aggregate scores. First, we evaluated how much of the variance of the multivariate intersubject similarity matrices could be accounted for by the intersubject distances of the summary scores. The spearman rho between the multivariate and univariate summary score distance matrices for sociosexual desire was, *r* = .30, *p* = .0008 (Figure 5A & 5C). We observed a similar relationship for the corresponding analysis for the self-control matrices, *r* = .30, *p* < .001 (Figure 5B & 5D). Second, we ran an IS-RSA on the summary score intersubject distance matrices and found that no regions survived multiple comparison correction (FDR) for either the sociosexual desire or self-control preferences (Figure S1). This suggests that a single univariate summary score captures at most 9% of the variance of its respective multivariate counterpart and that relying on the higher intrinsic variance of the multivariate distances is critical to adequately capturing variation in affective brain responses.

### Meta-analytic Decoding

Finally, we used meta-analytic decoding to identify which psychological constructs were most likely involved in constructing the affective experience given the pattern of brain results we observed from the un-thresholded IS-RSA maps. When participants were watching erotic movies, inter-subject similarity brain maps based on variations in sociosexual desire revealed stronger associations with the social and social judgment topic maps from Neurosynth, whereas variations in self-control revealed stronger associations with the emotion, reward, executive control, cognitive control, error monitoring, and conflict topic brain maps (Figure 6C).

## Discussion

In this study, we used brain imaging to explore the affective experience of heterosexual male participants. We used a naturalistic experimental design (Hasson et al., 2004; Haxby et al., 2011), in which participants watched short clips of erotic and neutral movies in the MRI scanner. These types of designs are ideal for eliciting powerful psychological experiences and creating strong variation in brain activity underlying the experience (Jolly and Chang, 2019). In contrast to the standard practices in emotion research, we did not examine affective experience using self-reported feelings or videos selected to elicit specific emotional states (Coan et al., 2007; Lench et al., 2011; Lindquist et al., 2012; Quigley et al., 2014). Instead, we explored how variation in two distinct preferences (i.e., sociosexual desires and self-control) mapped onto individual variation in brain dynamics while watching the videos using intersubject representational similarity analysis (IS-RSA).

Consistent with our predictions, we found that when individuals watched erotic movies, individuals with similar sociosexual desire preferences showed higher similarities in patterns of neural dynamics in brain regions within the cortico-striatal reward and default mode and mentalizing networks than those with different preferences. In contrast, as individuals became closer in their self-control preferences, we observed greater similarities in patterns of neural dynamics in brain regions within the fronto-parietal executive control and cingulo-insula salience networks. Importantly, when individuals watched neutral movies, inter-subject similarities in sociosexual desire and self-control preferences played no prominent role in accounting for the similarities in patterns of neural dynamics. We used meta-analytic decoding to provide a crude reverse inference of the possible psychological states contributing to the affective experience. Consistent with our expectations, variations in sociosexual desire preferences revealed stronger associations with social and social judgment topics, whereas variations in self-control preferences revealed stronger associations with the executive control, cognitive control, error monitoring and conflict. Together, our results support our hypothesis that variation in individual preferences can be used to explore affective experiences. Though we only specifically examined preferences for sociosexual desire and self-control, we do not believe this to be an exhaustive list of possible preferences and speculate that many other potential measures might also provide insight into this experience.

This study provides important conceptual and methodological advances to the investigation of affective experiences. Because there currently exists no objective measure of affective experiences (Chang et al., 2015), the field of emotion has a long history of grappling with measurement issues and has largely relied on self-report (Larsen and Fredrickson, 1999). One issue with trying to have participants map an experience into a high dimensional space of self-reported feelings is that this process requires both introspection (Nisbett and Wilson, 1977) and verbally labeling feelings using shared concepts (Lindquist et al., 2015). It’s possible that this verbal labeling process necessarily reduces the dimensionality of the representational space of the experience by filtering out processes that cannot be measured using this approach, which is why many studies find that 2-5 dimensions can explain the majority of the emotion rating variance (Chikazoe et al., 2014; Kragel and LaBar, 2015; Skerry and Saxe, 2015). In addition, most studies select a few stimuli to elicit a finite set of emotional states. However, this approach assumes that all participants will have a similar experience (Chang et al. 2018) and limits the variation in the emotional experiences (Cowen and Keltner, 2017), which can provide a statistical bias towards a low dimensional representation (Jolly and Chang, 2019). Our study provides an alternative approach to exploring affective experiences. Rather than assuming that participants will have the same response, which provides the basic premise of intersubject correlation (Hasson et al., 2004) and also functional alignment techniques (Guntupalli et al., 2016; Haxby et al., 2011), we assume that participants will have strong variations in their experience, which should correspond to structured variations in measures related to the experience. Importantly, we do not attempt to reduce the dimensionality of these measures to a single summary score, instead, we represent each item from the measure as a separate axis in a multiple-dimensional space and calculated the pairwise distance of each participant in this high-dimensional space. We believe that preserving the richness and complexity of all features in a high-dimensional space is important as there are many ways to answer a questionnaire that will produce an identical single summary score. Consistent with this intuition, we find that our IS-RSA results only hold when using a high dimensional representation and are not present when mapping participants’ distances using summary scores.

Though we believe this IS-RSA approach to be promising, there are several important limitations that should be acknowledged. First, we are using all of the features of each preference measure as an axis to map each participant and weighting the contribution of each feature equally. This means that we currently are unable to determine *which* features are specifically contributing to the experience. In addition, some of these features could be reflecting pure noise, which would be weighted equally as features that contain pure signal. It’s possible that this might be addressed with future work using multivariate regression techniques (e.g., partial least squares). Second, we are mapping individual position in this multidimensional space to intersubject dynamics of brain activity. This means that we do not know *when* in time processes specific to the experience occurred. It is possible to use similarity in spatial representations (van Baar et al., 2019), which might provide a way to extend this to which time points show a similar intersubject structure (Chang et al. 2018). However, this will also require accounting for multiple comparisons as well as non-independence in the time-series signals resulting from autocorrelation.

In summary, we have provided a demonstration of how variations in participants’ preferences can be used to uncover unique dimensions of an affective experience based on similarity in the intersubject structure of brain dynamics measured during the experience. This technique has the potential to provide a new approach to studying the neural processes underlying psychological experiences elicited through naturalistic experimental designs. Though this study provides a simple proof of concept, we hope that this work will inspire future innovations in analyzing naturalistic experimental designs, affective science, and psychological experiences.

## Acknowledgments

The authors wish to thank Ying Lin and Mary D’Geronimo for help with data collection. This work was supported by funding from the National Institute of Mental Health R01MH116026, R56MH080716, and National Institute on Drug Abuse R01DA022582. This work was also supported by funding from Chiang Ching-Kuo Foundation for International Scholarly Exchange (GS040-A-16) and from the Ministry of Science and Technology (MOST) in Taiwan (Young Scholar Fellowship Program, grant number: 108WXA0310050). Data and code used to perform the analysis in the paper will be available on github pending publication of this manuscript (https://github.com/cosanlab/affective_ISRSA).

